# The 2009 pandemic H1N1 hemagglutinin stalk remained antigenically stable after circulating in humans for a decade

**DOI:** 10.1101/2022.01.03.474869

**Authors:** Shannon R. Christensen, Emily T. Martin, Joshua G. Petrie, Arnold S. Monto, Scott E. Hensley

## Abstract

An H1N1 influenza virus caused a pandemic in 2009 and descendants of this virus continue to circulate seasonally in humans. Upon infection with the 2009 H1N1 pandemic strain (pH1N1), many humans produced antibodies against epitopes in the hemagglutinin (HA) stalk. HA stalk-focused antibody responses were common among pH1N1-infected individuals because HA stalk epitopes were conserved between the pH1N1 strain and previously circulating H1N1 strains. Here, we completed a series of experiments to determine if the pH1N1 HA stalk has acquired substitutions since 2009 that prevent the binding of human antibodies. We identified several amino acid substitutions that have accrued in the pH1N1 HA stalk from 2009-2019. We completed enzyme-linked immunosorbent assays, absorption-based binding assays, and surface plasmon resonance experiments to determine if these substitutions affect antibody binding. Using sera collected from 230 humans (aged 21-80 years), we found that pH1N1 HA stalk substitutions that have emerged since 2009 do not affect antibody binding. Our data suggest that the HA stalk domain of pH1N1 viruses remained antigenically stable after circulating in humans for a decade.

**Importance:** In 2009, a new pandemic H1N1 (pH1N1) virus began circulating in humans. Many individuals mounted hemagglutinin (HA) stalk-focused antibody responses upon infection with the 2009 pH1N1, since the HA stalk of this virus was relatively conserved with other seasonal H1N1 strains. Here, we completed a series of studies to determine if the 2009 pH1N1 strain has undergone antigenic drift in the HA stalk domain over the past decade. We found that serum antibodies from 230 humans could not antigenically distinguish the 2009 and 2019 HA stalk. These data suggest that the HA stalk of pH1N1 has remained antigenically stable, despite the presence of high levels of HA stalk antibodies within the human population.

## Introduction

Influenza pandemics can occur when a novel influenza strain crosses the species barrier into humans. Limited immunological memory within the population can allow for unhindered spread of the novel virus, resulting in high morbidity and mortality rates [1]. After the population develops robust immunity, pandemic influenza viruses can persist in the population by evading previously elicited antibody (Ab) responses through the accumulation of substitutions in the viral membrane proteins hemagglutinin (HA) and neuraminidase (NA), through a process called antigenic drift [2]. Amino acid substitutions in antigenic sites of HA can arise to directly prevent antibody binding [3], or can be stochastically selected by other factors to fine-tune HA function, such as altering pH of fusion [4] or viral binding to host receptors [5].

In 2009, a novel triple-reassortant H1N1 virus caused a pandemic [6]. The 2009 pandemic was unusual because most of the population already had pre-existing immunity to other H1N1 strains [7]. The 2009 pandemic H1N1 (pH1N1) HA protein possessed an antigenically distinct globular head domain relative to previously circulating H1N1 (sH1N1) strains [8-11], which allowed the virus to circumvent most pre-existing human neutralizing Abs. However, the HA stalk domain of the 2009 pH1N1 strain was relatively similar to the HA stalk domain of previously circulating sH1N1 viruses [8-10]. Upon infection with the 2009 pH1N1 virus, memory B cells specific for the HA stalk were preferentially recalled in many humans, leading to Ab responses that were highly focused on the HA stalk [12-14].

Human challenge studies showed that higher levels of HA stalk antibodies correlate with reduced viral shedding and number of symptoms upon pH1N1 infection [15], and serological cohort studies demonstrated that HA stalk antibodies can be associated with protection against pH1N1 influenza virus infection in adults and children [16, 17]. Some HA stalk-specific Abs neutralize antigenically distinct influenza viruses within and across influenza A group 1 and group 2 viruses [13, 18, 19], and a small number neutralize both influenza A and influenza B strains in mouse and ferret models [20, 21]. Given that HA stalk Abs are broadly reactive, several new vaccine approaches are being developed to elicit these Abs [22].

Although the HA stalk domain is typically more constrained than the HA globular head [23], it is possible that high levels of antibodies against the HA stalk of pH1N1 in the human population may drive antigenic drift within this domain. Indeed, the pH1N1 HA stalk domain is able to acquire substitutions *in vitro* when incubated with HA stalk-specific monoclonal (mAbs) or human polyclonal serum [24]. In these experiments, the isolated HA stalk escape mutants not only replicated well *in vitro*, but also retained pathogenicity *in vivo* [24]. Consistent with this, a recent pH1N1 human challenge study demonstrated that HA stalk mutants are enriched in individuals who have high levels of HA stalk antibodies [25]. Collectively, these studies challenge the dogma that the HA stalk domain is resistant to antigenic change.

Here, we completed experiments to determine if the HA stalk of pH1N1 acquired antigenic changes after circulating for 10 years in humans. We identified 7 HA stalk amino acid substitutions that have become fixed in pH1N1 viruses since 2009 and we characterized the antigenic effects of these substitutions using human antibodies.

## Results

We analyzed the HA sequences of 2,047 pH1N1 influenza genomes to identify amino acid substitutions that rose to fixation in the stalk domain from 2009-2019 [26, 27]. A total of seven amino acid substitutions were identified (**Fig. 1A**) that accrued over a span of ten years of seasonal circulation (**Fig. 1B**). Four amino acid substitutions are located in the HA1 domain of the HA stalk (A13T, K283E, I295V, and I324V) and three amino acid substitutions are located in the HA2 domain (E47K, S124N, and E172K) (**Fig. 1A**). To determine if these substitutions affect human antibody binding, we created a ‘headless’ 2019 H1 recombinant protein for serological analyses. To do this, we introduced five of the substitutions (HA1-A13T, HA1-I324V, HA2-E47K, HA2-S124N, and HA2-E172K) into a 2009 pH1N1 headless HA stalk recombinant protein. We did not include the HA1-K283E and HA1-I295V substitutions since these amino acids are located closer to the HA head outside the area included in our headless HA construct.

**FIG. 1.**
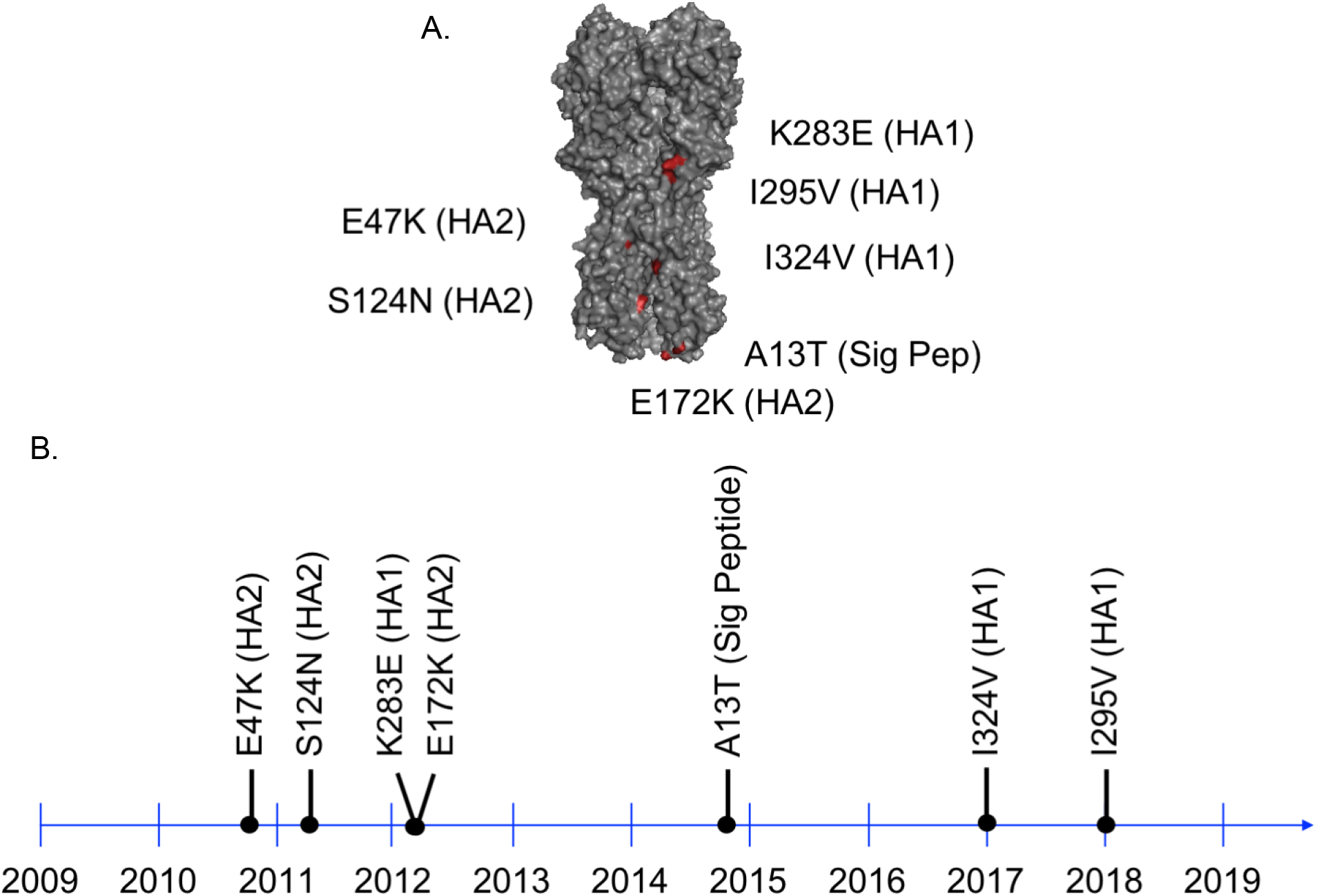
Amino acid substitutions have accrued in the HA stalk domain. We analyzed amino acid positions in 2,047 pH1N1 HA genomes from 2009 - 2019 using Nextstrain.org. (**A**) Seven amino acid substitutions emerged in the seasonal H1N1 HA stalk domain from 2009-2019. (**B**) Most of the pH1N1 HA stalk substitutions emerged after the 2011-2012 influenza season.

We used enzyme-linked immunosorbent assays (ELISAs) to determine if human antibodies differentially recognized the 2009 pH1N1 HA headless stalk protein (abbreviated as 2009 HA stalk) or the 2009 pH1N1 HA headless stalk protein containing the five amino acid substitutions (abbreviated as 2019 HA stalk). For these experiments, we analyzed serum samples collected from 230 participants enrolled in an Ann Arbor, Michigan household cohort study during the 2011-2012 influenza season. We analyzed samples from the 2011-2012 season because many individuals were likely previously infected or vaccinated with pH1N1 at that point but most of the HA stalk substitutions emerged after the 2011-2012 season. Serum antibodies from most participants reacted similarly to the 2009 and 2019 HA stalk constructs, with a relative 2009 stalk mean titer of 3052 and 2019 stalk mean titer of 2560 (**Fig. 2A**). Only 1 of 230 samples tested had >2-fold difference in antibody titers when tested against the 2009 and 2019 H1 HA stalks (**Fig. 2B**). Antibody titers against the 2009 and 2019 H1 HA stalks were highly correlated (**Fig. 2C**).

**FIG. 2.**
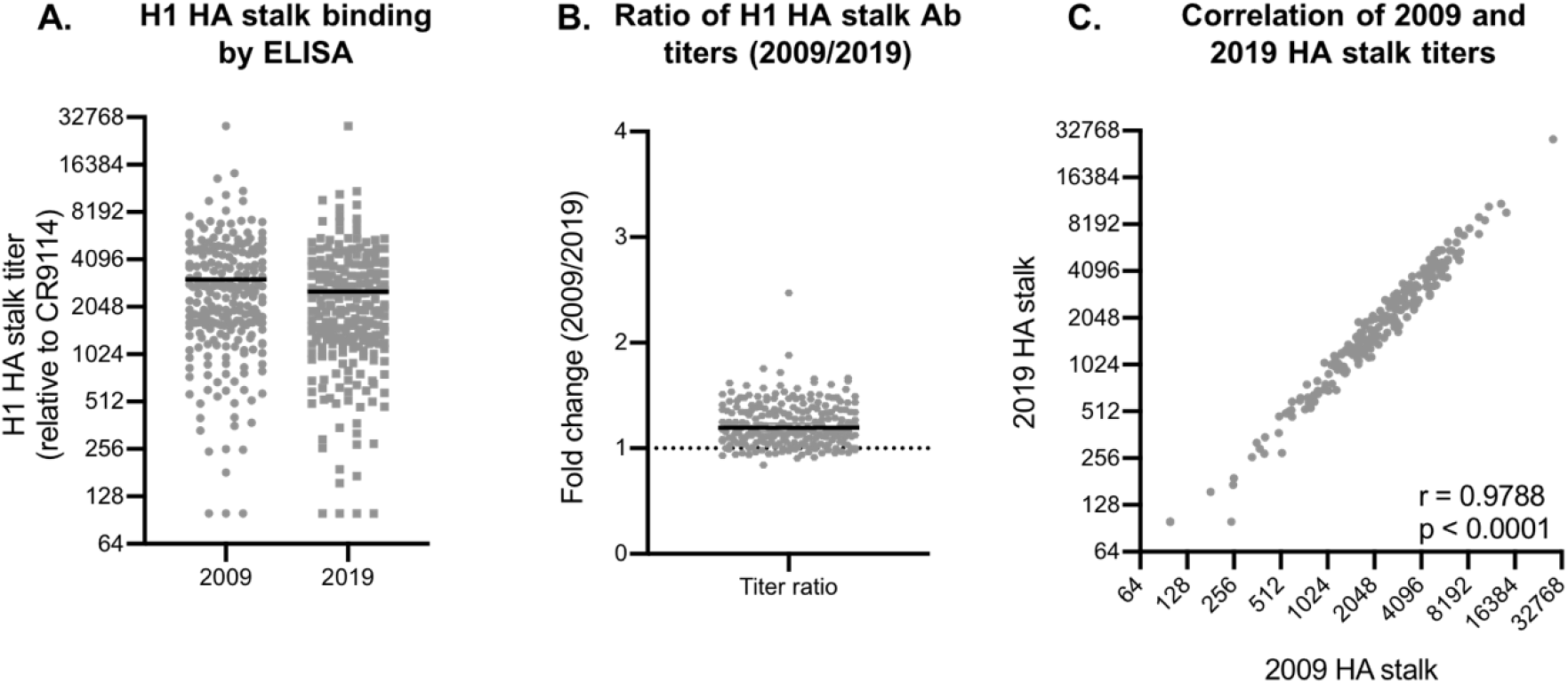
Amino acid substitutions do not greatly reduce binding of human polyclonal antibodies. (**A**) ELISAs were used to quantify 2009 and 2019 HA stalk antibody titers in human serum samples collected before the 2011-2012 influenza season. Each dot represents antibody titers of a single serum sample (n = 230). Relative mean stalk titers are 3,052 and 2,560 for the 2009 and 2019 H1 HA stalk domain, respectively. (**B**) 229 samples out of 230 demonstrated <2-fold difference in titers between the 2009 and 2019 H1 HA stalks. The ratio of the titers (2009/2019) were determined using geometric mean titers from two independent experiments. (**C**) Relative titers between the 2009 and 2019 HA stalk were strongly and significantly correlated (Spearman correlation, r = 0.9788, p < 0.0001).

We next performed surface plasmon resonance (SPR) with a subset of serum samples to determine if the amino acid substitutions in the 2019 H1 stalk reduced binding affinity of serum antibodies from the 2011-2012 household participants. Representative curves and associated relative Kd values are depicted in **Fig. 3A** and **Table 1**. Consistent with our ELISA data, antibodies from each sample had similar Kd values when assays were completed with the 2009 and 2019 HA stalk (**Fig. 3B** and **Table 1**) and 2009 and 2019 Kd values were highly correlated (**Fig. 3C**).

**Table 1.** Relative Kd values for H1 HA stalk-specific antibodies in human polyclonal serum. Each serum sample was assessed by surface plasmon resonance (SPR) against the 2009 and 2019 headless HA stalk. Relative Kd values represent the geometric mean value of two independent experiments.

| Sample ID | 2009 H1 HA stalk Kd* | 2019 H1 HA stalk Kd* |
| --- | --- | --- |
| H0029 | 8.86e-11 | 8.39e-11 |
| H0050 | 3.48e-10 | 3.39e-10 |
| H0054 | 5.87E-10 | 5.45E-10 |
| H0070 | 2.40E-09 | 2.22E-09 |
| H0083 | 3.08e-10 | 2.37e-10 |
| H0098 | 5.41e-10 | 4.46e-10 |
| H0152 | 1.29e-9 | 1.19e-9 |
| H0156 | 3.62e-11 | 3.02e-11 |
| H0166 | 2.00e-9 | 1.81e-9 |
| H0181 | 1.05e-9 | 8.86e-10 |
| H0182 | 1.85e-10 | 1.63e-10 |
| H0191 | 3.68e-7 | 3.05e-7 |
| H0194 | 1.09E-06 | 1.05E-06 |
| H0197 | 4.43e-8 | 3.81e-8 |
| H0216 | 7.90e-7 | 7.12e07 |
| H0219 | 2.98E-08 | 2.48E-08 |

\*Relative to 20 nM CR9114

**FIG. 3.**
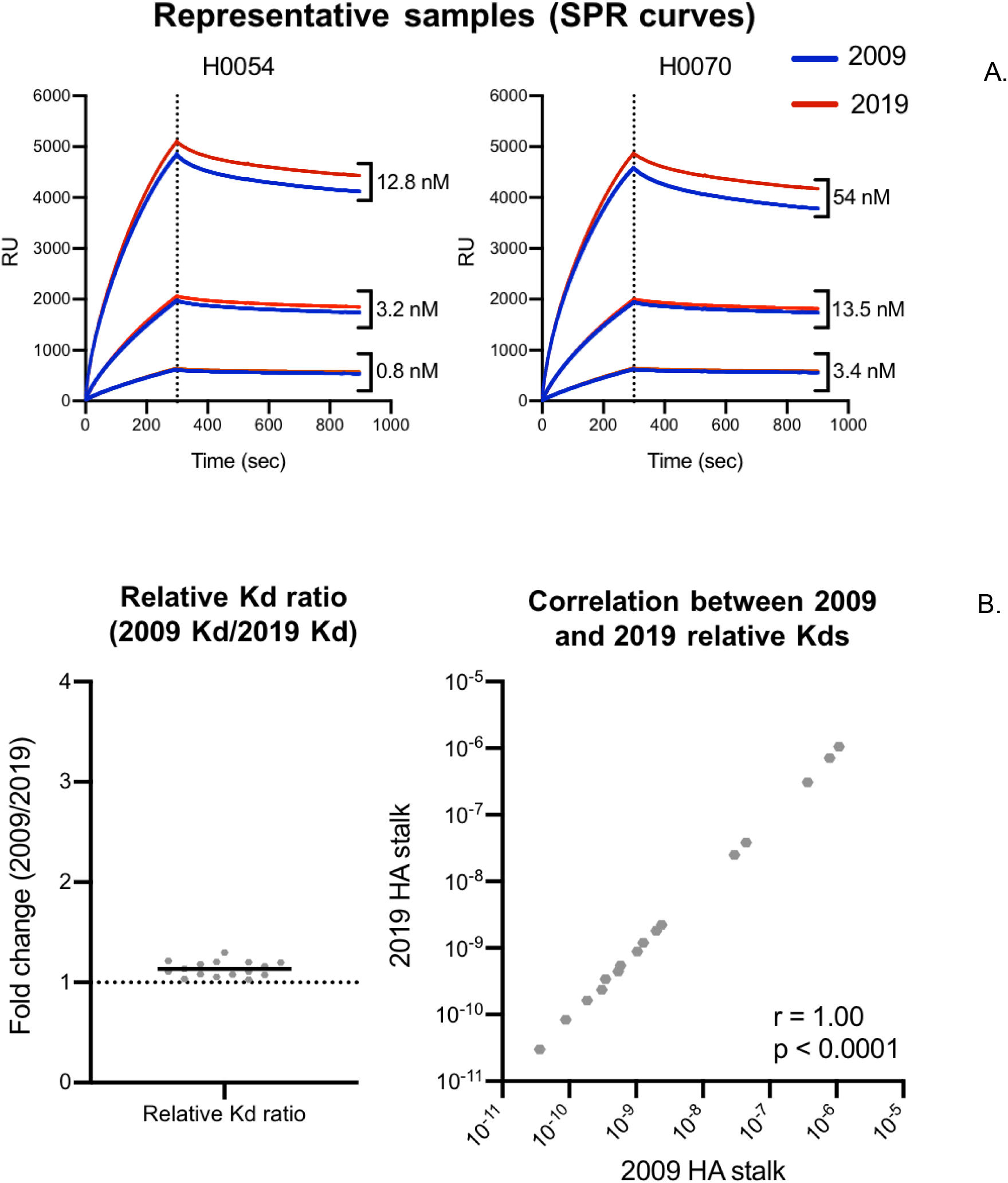
Amino acid substitutions do not decrease affinity of H1 HA stalk-specific antibodies in human polyclonal serum. A subset of serum samples (n = 16) were analyzed for relative Kd values against the 2009 and 2019 headless HA stalk by surface plasmon resonance (SPR). (**A**) Representative serum sample binding and dissociation curves by SPR are showed for 2 serum samples (blue line = 2009 stalk, red line = 2019 stalk). (**B**) All samples demonstrated <2-fold difference in relative Kd values between the 2009 and 2019 H1 HA stalks. The ratios of the titers (2009/2019) were determined using geometric mean Kd values of two independent experiments. Each dot represents values from a single serum sample. (**C**) Relative Kd values for the 2009 and 2019 stalks were highly correlated (Spearman correlation, r2 = 1.00, p < 0.0001). Correlation was performed on geometric mean titers of two independent experiments. Each dot represents values from a single serum sample.

Finally, we completed absorption assays to verify that human serum antibodies bound to both the 2009 and 2019 HA stalks. For these experiments, we transfected cells with constructs expressing the 2009 or 2019 HA stalk, and then we incubated a subset of serum samples with these cells to absorb out HA stalk-reactive Abs. We then completed ELISAs with the absorbed serum samples to quantify antibodies that bound to the 2009 and 2019 HA stalks. In these assays, antibodies that bind to the 2009 HA stalk but not to the 2019 HA stalk can be absorbed with cells expressing the 2009 HA stalk but not the 2019 HA stalk. However, we did not observe these types of antibodies in human serum samples. We found that both the 2009 and 2019 HA stalk efficiently absorbed all HA stalk-specific antibodies from each serum sample (**Fig. 4**), indicating that antibodies reactive to the 2009 HA stalk bind efficiently to the 2019 HA stalk.

**FIG. 4.**
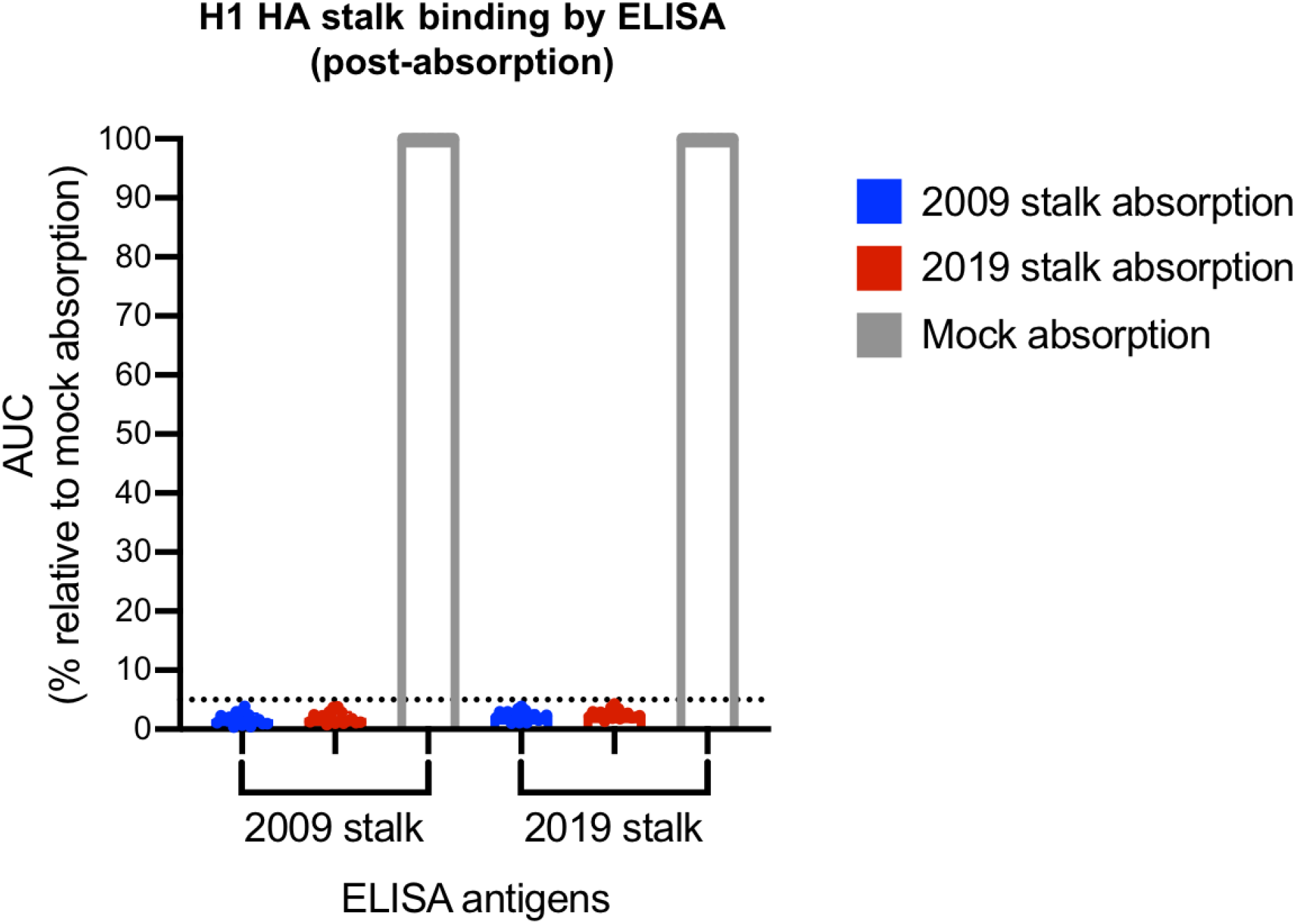
Amino acid substitutions in the H1 HA stalk domain do not decrease absorption efficiency of human polyclonal serum antibodies. Serum samples were absorbed with cells expressing either the 2009 or 2019 HA stalk. HA stalk-reactive antibodies remaining after absorptions were quantified by ELISA. HA stalk titers are depicted as % area under the curve (AUC) relative to the mock absorption condition. Absorption with cells expressing either the 2009 or 2019 HA stalk completely removed antibodies that bound to either HA, confirming that these HA stalks are antigenically similar.

## Discussion

Antigenic drift in the HA globular head has been relatively well-studied [3, 5, 28-30]. However, much less is known about evolution within the HA stalk domain, despite this being a major target for many universal influenza vaccine strategies. There is evidence of positive selection of amino acid substitutions in the HA stalk domain pH1N1 viruses [31], and *in vitro* work has demonstrated that H1 HA stalk escape mutants can be selected for when pH1N1 is cultured in the presence of human immune sera or stalk-specific mAbs [24]. In this report we show that pH1N1 viruses have acquired several substitutions in the HA stalk since 2009, but these substitutions do not greatly affect human antibody binding.

It is possible that the HA stalk substitutions that have emerged since 2009 were selected as a means to improve HA function. One of the HA stalk amino acid substitutions that appeared early on, E47K, has been closely studied for its effect on viral function [4]. The E47K substitution increases both acid and thermal stability *in vitro*, and also increases pathogenicity *in vivo* [4]. While mechanistic studies have not been conducted on the other amino acid substitutions that have arisen in the pH1N1 HA stalk domain since 2009, it is possible they also contribute to fine-tuning of HA function.

It is also possible that substitutions in the pH1N1 HA stalk arose to compensate for antigenic changes within the HA head domain, a mechanism of viral evolution that has previously been identified in the highly pathogenic avian influenza virus subtype H5N1 [32]. Amino acid substitutions that mediate antigenic escape often incur pleiotropic effects, such as altered binding to sialic acid, the host cellular receptor for influenza viruses, or decreased protein-folding stability [33-38], which in turn decrease viral fitness. In many cases viral fitness can be restored by compensatory mutations in the HA or NA proteins that have opposing effects [34-36, 38-41]. Many compensatory mutations are neutral and will arise through random drift first [42], thus allowing an otherwise deleterious antigenic mutation to arise, or the two mutations may occur simultaneously. As such, the evolution of compensatory mutations can lead to entrenchment, the inability of a substitution to revert to its ancestral state without deleterious effects [43].

While the findings presented in this study do not negate the potential for antigenic drift to occur within the pH1N1 HA stalk domain, they do support the HA stalk domain as a durable target for a universal influenza vaccine. Future studies should focus on the durability of HA stalk domain immune responses after immunization with HA stalk vaccine platforms, as well as the elicitation of these responses in humans with different viral exposure histories.

## Materials and Methods

### Human subjects

During the 2011-2012 influenza season, adult (≥18 years) participants were prospectively enrolled in an observational, household cohort design study of influenza vaccine effectiveness [44]. All participants provided informed consent and completed an enrollment interview. Influenza vaccination status was defined by self-report and documentation in the electronic medical record and Michigan Care Improvement Registry (MCIR). All specimens were collected prior to the start of the influenza season. Studies involving humans were approved by the Institutional Review Boards of the University of Michigan and University of Pennsylvania. All experiments (ELISAs, absorptions, and surface plasmon resonance) were completed at the University of Pennsylvania using deidentified sera.

### Recombinant HA proteins

Plasmids encoding the 2009 H1 HA stalk were provided by Adrian McDermott and Barney Graham from the Vaccine Research Center at the National Institutes of Health. We created plasmids to express the 2019 H1 HA stalk that incorporated five amino acid substitutions (HA1-A13T, HA1-I324V, HA2-E47K, HA2-S124N, and HA2-E172K). We also created plasmids to express the 2009 and 2019 H1 HA stalks with a transmembrane domain for absorption studies. We created pSport6-based plasmids after obtaining gBlock genes from Integrated DNA Technologies. Headless HA stalk proteins were expressed in 293F cells and purified using nickel-nitrilotriacetic acid agarose (no. 1018244, Qiagen) in 5-ml polypropylene columns (no. 34964; Qiagen), washed with 50 mM Na_2_HCO_3_, 300 mM NaCl, and 20 mM imidazole buffer at pH 8. The protein was then eluted in 50 mM Na_2_HCO_3_, 300 mM NaCl, and 300 mM imidazole buffer at pH 8. Purified protein was then buffer exchanged into phosphate-buffered saline (PBS; no. 21-031-CM; Corning). H1 HA stalk proteins were aliquoted and stored at -80C. For some experiments, H1 HA stalk proteins were biotinylated using the Avidity BirA-500 kit (no. BirA500) and stored in aliquots at -80C.

### mAbs

Plasmids encoding the human mAb CR9114 were provided by Patrick Wilson (University of Chicago). CR9114 was expressed in 293T cells and purified 4 days post-infection using NAB protein A/G spin kits (no. 89950; Thermo Fisher).

### Headless HA ELISAs

Headless HA ELISAs were performed using 96-well Immunlon 4HBX flat-bottom microtiter plates (no. 3855; Thermo Fisher) coated with 0.5 ug/well of streptavidin (no. S4762; Sigma). We completed total IgG headless HA ELISAs (using both the 2009 and 2019 stalks) with all serum samples. A detailed protocol has been previously described elsewhere [16]. In brief, headless HA stalk proteins were diluted in biotinylation buffer to a concentration of 0.25 ug/ml and 50 ul were added to each well. Wells were blocked with 150 ul of biotinylation blocking buffer. Serum samples were serially diluted in biotinylation buffer (starting at 1:100 diltiion), then added to the ELISA plates. The human, HA stalk-specific mAb CR9114 was used as a plate control, starting at 0.03 ug/ml, to verify equal coating of plates and to determine relative serum titers. Peroxidase-conjugated goat anti-human IgG (no. 109-036-098; Jackson) was added at 50 ul/well. Finally, SureBlue TMB peroxidase substrate (no. 5120-0077; KPL) was added to each well and the reaction was quenched with the addition of 25 ul/well of 250 mM HCl solution. Each step was incubated for an hour at room temperature on a rocker. Plates were extensively washed with PBS (no. 21-031-CM) and 0.1% Tween 20 between each step using a BioTek 405 LS microplate washer. Relative titers were determined based on CR9114 mAb binding for each plate, and reported as the corresponding inverse of the serum dilution that generated the equivalent optical densities (OD). Each ELISA was performed a minimum of two times.

### Absorption ELISAs

Absorption ELISAs were completed using a subset of serum samples. 293F cells were split to a 1e6 cells/ml density in 500 mL (160 ml cell culture volume) vented tissue culture flasks (no. 431145; Corning) on the day of transfection. 293F cells were transfected using 1 ml Opti-MEM (no. 31985-070; Gibco), 320 ul 293Fectin (no. 12347-019; Gibco), and 160 ug plasmids containing the H1 HA stalk constructs with a transmembrane domain, or a mock transfection was performed. The cells were incubated at 37C, shaking at 800 rpm, for 3 days. The cells from each transfection condition were collected, spun down at 300 xg for 5 minutes, then re-suspended in 30 mls FreeStyle 293 Expression Medium (no. 12338018; Thermo Fisher). An aliquot of cells was taken for counting and the cells were spun again at 300 xg for 5 minutes, then re-suspended in an appropriate volume of FreeStyle 293 Expression Medium (no. 12338018; Thermo Fisher) to obtain a concentration of 1e8 cells/ml. Cells and individual serum samples at a 1:10 dilution were mixed in a 1:1.1 ratio (to accommodate for the substantial volume of cells) to ensure an excess of antigen was present. The serum/cell mixtures were incubated in 96-well U-bottom plates (no. 353077; Corning) for 1 hour on a plate shaker set to 800 rpm. The 96-well plates were then centrifuged at 2000 RPM for 5 minutes and the supernatants transferred to different 96-well U-bottom plates (no. 353077; Corning) and centrifuged a second time for 5 minutes at 2000 rpm to remove all cells. The supernatants were then tested by ELISA for HA stalk-reactive antibodies. As an absorption control, the human mAb CR9114 was incubated under the same absorption conditions at a concentration of 2 ug/ml. Each absorption ELISA was performed a minimum of two times.

### Surface plasmon resonance

All SPR experiments were performed on the BIACore 3000 instrument by GE Healthcare. The LS Sensor Chip NTA (no. BR-1004-07; GE Health Sciences) surface was conditioned by injecting 350 mM EDTA (diluted from 0.5 M, pH 8 stock, no. 15575020; Thermo Fisher) at 10 ul/min for 1 minute followed by extensive washing with running buffer to remove residual EDTA. To ensure all EDTA was removed, a subsequent washing step using 50 mM NaOH, injected at 10 ul/min for 1 minute was performed. The chip surface was then prepared for antigen capture by injecting 0.5M NiCl2 at 10 ul/min for 1 minute. The HIS-tagged 2009 and 2019 HA stalk proteins, previously diluted to 2.5 ug/ml in 1x HBS-P buffer (0.01 M HEPES (no. 25-060-Cl; Corning) pH 7.4, 0.15 M NaCl, 0.005% v/v Polysorbate 20 (Tween 20) (no. BP337-100; Thermo Fisher)), were immobilized onto separate channels of the LS Sensor Chip NTA (no. BR-1004-07; GE Health Sciences) at 5 ul/min with a target density of 350 response units, followed by a 20 minute incubation. Serum samples, previously diluted to concentrations of 1:100, 1:400, and 1:1600 in 1x HBS-P buffer, were injected at 20 ul/min for 150 seconds over the immobilized rHA stalk chip channels or the reference chip channel, followed by a 400s dissociation phase. Each step was followed by an extensive washing cycle using 1x HBS-P buffer. For kinetic analysis, values obtained after injections over reference cell channels and injections with buffer only were subtracted from the data. Association rates (k_a_), dissociation rates (k_d_), and equilibrium dissociation constants (K_d_) were calculated by aligning the curves to fit a Langmuir 1:1 binding model using BIAevaluation 4.1 software. To obtain a relative concentration of H1 HA stalk-specific antibodies in each polyclonal serum sample used, we performed an HA stalk ELISA (according to methods described above) to determine the serum dilution required to achieve an OD equivalent to 2 nM of CR9114. These relative serum concentrations were used as input for the Langmuir 1:1 model for each serum sample concentration. Each SPR experiment was performed in triplicate.

## Acknowledgments

This project has been funded in part with Federal funds from the National Institute of Allergy and Infectious Diseases, National Institutes of Health, Department of Health and Human Services, under Contract No. 75N93021C00015, Contract No. HHSN272201, and Grant No. 1R01AI108686. SEH holds an Investigators in the Pathogenesis of Infectious Disease Awards from the Burroughs Wellcome Fund.

## References

1. Cox, N.J. and K. Subbarao, Global epidemiology of influenza: past and present. Annu Rev Med, 2000. 51: p. 407–21.

2. CDC. How the flu virus can change: “drift” and “shift”. 2019 11/15/2019]; Available from: https://www.cdc.gov/flu/about/viruses/change.htm.

3. Caton, A.J., et al., The antigenic structure of the influenza virus A/PR/8/34 hemagglutinin (H1 subtype). Cell, 1982. 31(2 Pt 1): p. 417–27.

4. Cotter, C.R., H. Jin, and Z. Chen, A single amino acid in the stalk region of the H1N1pdm influenza virus HA protein affects viral fusion, stability and infectivity. PLoS Pathog, 2014. 10(1): p. e1003831.

5. Hensley, S.E., et al., Hemagglutinin receptor binding avidity drives influenza A virus antigenic drift. Science, 2009. 326(5953): p. 734–6.

6. Smith, G.J., et al., Origins and evolutionary genomics of the 2009 swine-origin H1N1 influenza A epidemic. Nature, 2009. 459(7250): p. 1122–5.

7. Cobey, S. and S.E. Hensley, Immune history and influenza virus susceptibility. Curr Opin Virol, 2017. 22: p. 105–111.

8. Garten, R.J., et al., Antigenic and genetic characteristics of swine-origin 2009 A(H1N1) influenza viruses circulating in humans. Science, 2009. 325(5937): p. 197–201.

9. Manicassamy, B., et al., Protection of mice against lethal challenge with 2009 H1N1 influenza A virus by 1918-like and classical swine H1N1 based vaccines. PLoS Pathog, 2010. 6(1): p. e1000745.

10. Skountzou, I., et al., Immunity to pre-1950 H1N1 influenza viruses confers cross-protection against the pandemic swine-origin 2009 A (H1N1) influenza virus. J Immunol, 2010. 185(3): p. 1642–9.

11. Xu, R., et al., Structural basis of preexisting immunity to the 2009 H1N1 pandemic influenza virus. Science, 2010. 328(5976): p. 357–60.

12. Pica, N., et al., Hemagglutinin stalk antibodies elicited by the 2009 pandemic influenza virus as a mechanism for the extinction of seasonal H1N1 viruses. Proc Natl Acad Sci U S A, 2012. 109(7): p. 2573–8.

13. Wrammert, J., et al., Broadly cross-reactive antibodies dominate the human B cell response against 2009 pandemic H1N1 influenza virus infection. J Exp Med, 2011. 208(1): p. 181–93.

14. Thomson, C.A., et al., Pandemic H1N1 Influenza Infection and Vaccination in Humans Induces Cross-Protective Antibodies that Target the Hemagglutinin Stem. Front Immunol, 2012. 3: p. 87.

15. Park, J.K., et al., Evaluation of Preexisting Anti-Hemagglutinin Stalk Antibody as a Correlate of Protection in a Healthy Volunteer Challenge with Influenza A/H1N1pdm Virus. mBio, 2018. 9(1).

16. Christensen, S.R., et al., Assessing the Protective Potential of H1N1 Influenza Virus Hemagglutinin Head and Stalk Antibodies in Humans. J Virol, 2019. 93(8).

17. Ng, S., et al., Novel correlates of protection against pandemic H1N1 influenza A virus infection. Nat Med, 2019. 25(6): p. 962–967.

18. Li, G.M., et al., Pandemic H1N1 influenza vaccine induces a recall response in humans that favors broadly cross-reactive memory B cells. Proc Natl Acad Sci U S A, 2012. 109(23): p. 9047–52.

19. Qiu, C., et al., Boosting heterosubtypic neutralization antibodies in recipients of 2009 pandemic H1N1 influenza vaccine. Clin Infect Dis, 2012. 54(1): p. 17–24.

20. Dreyfus, C., et al., Highly conserved protective epitopes on influenza B viruses. Science, 2012. 337(6100): p. 1343–8.

21. Ekiert, D.C., et al., Antibody recognition of a highly conserved influenza virus epitope. Science, 2009. 324(5924): p. 246–51.

22. Erbelding, E.J., et al., A Universal Influenza Vaccine: The Strategic Plan for the National Institute of Allergy and Infectious Diseases. J Infect Dis, 2018. 218(3): p. 347–354.

23. Kirkpatrick, E., et al., The influenza virus hemagglutinin head evolves faster than the stalk domain. Sci Rep, 2018. 8(1): p. 10432.

24. Anderson, C.S., et al., Natural and directed antigenic drift of the H1 influenza virus hemagglutinin stalk domain. Sci Rep, 2017. 7(1): p. 14614.

25. Park, J.K., et al., Pre-existing immunity to influenza virus hemagglutinin stalk might drive selection for antibody-escape mutant viruses in a human challenge model. Nat Med, 2020. 26(8): p. 1240–1246.

26. Hadfield, J., et al., Nextstrain: real-time tracking of pathogen evolution. Bioinformatics, 2018. 34(23): p. 4121–4123.

27. Jover Lee, R.N., Trevor Bedford. Real-time tracking of influenza A/H1N1pdm evolution using data from GISAID. 2019 05/2019]; Available from: https://nextstrain.org/flu/seasonal/h1n1pdm/ha/12y.

28. Meyer, A.G. and C.O. Wilke, Geometric Constraints Dominate the Antigenic Evolution of Influenza H3N2 Hemagglutinin. PLoS Pathog, 2015. 11(5): p. e1004940.

29. Doud, M.B. and J.D. Bloom, Accurate Measurement of the Effects of All Amino-Acid Mutations on Influenza Hemagglutinin. Viruses, 2016. 8(6).

30. Thyagarajan, B. and J.D. Bloom, The inherent mutational tolerance and antigenic evolvability of influenza hemagglutinin. Elife, 2014. 3.

31. Su, Y.C.F., et al., Phylodynamics of H1N1/2009 influenza reveals the transition from host adaptation to immune-driven selection. Nat Commun, 2015. 6: p. 7952.

32. Hanson, A., et al., Identification of Stabilizing Mutations in an H5 Hemagglutinin Influenza Virus Protein. J Virol, 2015. 90(6): p. 2981–92.

33. Wu, N.C., et al., Diversity of Functionally Permissive Sequences in the Receptor-Binding Site of Influenza Hemagglutinin. Cell Host Microbe, 2017. 22(2): p. 247–248.

34. Das, S.R., et al., Fitness costs limit influenza A virus hemagglutinin glycosylation as an immune evasion strategy. Proc Natl Acad Sci U S A, 2011. 108(51): p. E1417–22.

35. Mitnaul, L.J., et al., Balanced hemagglutinin and neuraminidase activities are critical for efficient replication of influenza A virus. J Virol, 2000. 74(13): p. 6015–20.

36. Myers, J.L., et al., Compensatory hemagglutinin mutations alter antigenic properties of influenza viruses. J Virol, 2013. 87(20): p. 11168–72.

37. Underwood, P.A., J.J. Skehel, and D.C. Wiley, Receptor-binding characteristics of monoclonal antibody-selected antigenic variants of influenza virus. J Virol, 1987. 61(1): p. 206–8.

38. Kosik, I., et al., Influenza A virus hemagglutinin glycosylation compensates for antibody escape fitness costs. PLoS Pathog, 2018. 14(1): p. e1006796.

39. Wu, N.C., et al., Diversity of Functionally Permissive Sequences in the Receptor-Binding Site of Influenza Hemagglutinin. Cell Host Microbe, 2017. 21(6): p. 742–753 e8.

40. Kryazhimskiy, S., et al., Prevalence of epistasis in the evolution of influenza A surface proteins. PLoS Genet, 2011. 7(2): p. e1001301.

41. Hensley, S.E., et al., Influenza A virus hemagglutinin antibody escape promotes neuraminidase antigenic variation and drug resistance. PLoS One, 2011. 6(2): p. e15190.

42. Weinreich, D.M., R.A. Watson, and L. Chao, Perspective: Sign epistasis and genetic constraint on evolutionary trajectories. Evolution, 2005. 59(6): p. 1165–74.

43. Wu, N.C., et al., A complex epistatic network limits the mutational reversibility in the influenza hemagglutinin receptor-binding site. Nat Commun, 2018. 9(1): p. 1264.

44. Ohmit, S.E., et al., Influenza vaccine effectiveness in the 2011-2012 season: protection against each circulating virus and the effect of prior vaccination on estimates. Clin Infect Dis, 2014. 58(3): p. 319–27.

